# Dysregulation of innate immune signaling in animal models of Spinal Muscular Atrophy

**DOI:** 10.1101/2023.12.14.571739

**Authors:** Eric L. Garcia, Rebecca E. Steiner, Amanda C. Raimer, Laura E. Herring, A. Gregory Matera, Ashlyn M. Spring

## Abstract

**Background:** Spinal Muscular Atrophy (SMA) is a devastating neuromuscular disease caused by hypomorphic loss of function in the Survival Motor Neuron (SMN) protein. SMA presents across broad spectrum of disease severity. Unfortunately, vertebrate models of intermediate SMA have been difficult to generate and are thus unable to address key aspects of disease etiology. To address these issues, we developed a *Drosophila* model system that recapitulates the full range of SMA severity, allowing studies of pre-onset biology as well as late-stage disease processes.

**Results:** Here, we carried out transcriptomic and proteomic profiling of mild and intermediate *Drosophila* models of SMA to elucidate molecules and pathways that contribute to the disease. Using this approach, we elaborated a role for the SMN complex in the regulation of innate immune signaling. We find that mutation or tissue-specific depletion of SMN induces hyperactivation of the Immune Deficiency (IMD) and Toll pathways, leading to overexpression of antimicrobial peptides (AMPs) and ectopic formation of melanotic masses in the absence of an external challenge. Furthermore, knockdown of downstream targets of these signaling pathways reduced melanotic mass formation caused by SMN loss. Importantly, we identify SMN as a negative regulator of an ubiquitylation complex that includes Traf6, Bendless and Diap2, and plays a pivotal role in several signaling networks.

**Conclusions:** In alignment with recent research on other neurodegenerative diseases, these findings suggest that hyperactivation of innate immunity contributes to SMA pathology. This work not only provides compelling evidence that hyperactive innate immune signaling is a primary effect of SMN depletion, but it also suggests that the SMN complex plays a regulatory role in this process *in vivo*. In summary, immune dysfunction in SMA is a consequence of reduced SMN levels and is driven by cellular and molecular mechanisms that are conserved between insects and mammals.

## BACKGROUND

Spinal Muscular Atrophy (SMA) is a neuromuscular disease caused by mutations in the human *Survival Motor Neuron 1* (*SMN1*) gene and the accompanying reduction in levels of SMN protein (Lefebvre *et al*. 1995). In humans and SMA animal models, complete loss of SMN function does not lead to SMA; it causes developmental arrest and early lethality (O’Hern et al. 2017). Hypomorphic point mutations in *SMN1* and/or reduced levels of full-length SMN protein cause the disease (Lefebvre *et al*. 1995, 1997). The age-of-onset and severity of the disease varies widely, leading to a historical classification of SMA into three distinct subtypes, Type I (Werdnig-Hoffman disease, early infantile onset), Type II (intermediate late infant onset), and Type III (Kugelberg-Welander, childhood onset) (Kugelberg and Welander 1956; Darras and Finkel 2017; Oskoui *et al*. 2017). More recently, clinicians have increasingly recognized that SMA is better characterized as a broad-spectrum disorder, ranging from severe (prenatal onset) to nearly asymptomatic (Dubowitz 2017; Singh *et al*. 2021). SMA phenotypic severity is inversely proportional to SMN protein levels; however, the proximal trigger of the disease remains a mystery.

Mouse models of intermediate or late-onset SMA have been difficult to generate. Mutations at the endogenous mouse *Smn* locus or copy number changes in human *SMN2* (an *SMN1* paralog) transgenes cause dramatic shifts in phenotype from mild and largely unaffected, to very severe, with onset of symptoms in utero and death between 4-14 days (reviewed in Burghes *et al*. 2017; Oskoui *et al*. 2017). To circumvent these problems, we developed a *Drosophila* model system (Praveen *et al*. 2012, 2014). Using a series of SMA-causing missense alleles, we have shown that this system recapitulates the wide-spectrum of phenotypic severity seen in human patients (Praveen *et al*. 2014; Garcia *et al*. 2016; Gray *et al*. 2018; Spring *et al*. 2019; Raimer *et al*. 2020; Gupta *et al*. 2021). Importantly, this system provides an opportunity to study all stages of the disease, from pre-onset biology to late-stage processes (Spring *et al*. 2019; Raimer *et al*. 2020; Gupta *et al*. 2021).

The phenotypes associated with *Drosophila* models of SMA include impaired locomotion, neuromuscular abnormalities, developmental delays, decreased viability, and reduced life span (Chan *et al*. 2003; Rajendra *et al*. 2007; Chang *et al*. 2008; Praveen *et al*. 2012, 2014; Imlach *et al*. 2012; Garcia *et al*. 2013; Spring *et al*. 2019). In notable agreement with the onset of the human disease, our fruitfly models of SMA also exhibit progressive loss of limb motility, displaying a more rapid decline in posterior versus anterior appendages (Spring *et al*. 2019). Additionally, specific mutations that affect the SMN Tudor domain were recently shown to affect SMN protein levels in a temperature sensitive manner (Raimer *et al*. 2020). Hence, *Drosophila* models of SMA are continuing to reveal how individual mutations disrupt SMN function, contributing to different aspects of the disease.

SMN protein is involved in the biogenesis of small nuclear ribonucleoproteins (snRNPs), core components of the spliceosome (Matera and Wang 2014). SMN carries out its functions in the assembly of snRNPs primarily in the cytoplasm (Matera and Wang 2014). *Smn* and *Phax (Phosphorylated Adaptor for RNA export)* null mutants exhibit an overlapping set of alternative splicing differences relative to wild-type animals (Garcia *et al*. 2016). Phax exports small nuclear RNAs (snRNAs) from the nucleus for assembly into snRNPs by Smn and the SMN complex (Ohno *et al*. 2000; Matera and Wang 2014). Recently, a common allele-specific *RpS21* alternative splicing event was shown to modify the larval lethality of *Phax*, but not *Smn*, mutants (Garcia 2022). Transcriptomic profiling of various *Smn* null and missense mutants has revealed the activation of an innate immune response that correlates with phenotypic severity of the different mutants (Garcia *et al*. 2013, 2016). Conspicuously, mutations in the *Phax* gene do not cause similar transcriptomic signatures of activated innate-immune signaling, which suggests that SMN may have a specific function in cellular immunity (Garcia *et al*. 2016).

Defects in the development of immune cells and tissues have been reported in several mouse models of SMA (Deguise and Kothary 2017; Khairallah *et al*. 2017; Thomson *et al*. 2017; Deguise *et al*. 2017). SMA model mice have smaller spleens and display altered red pulp macrophage morphology; events that reportedly precede evidence of neurodegeneration (Deguise and Kothary 2017; Khairallah *et al*. 2017; Thomson *et al*. 2017; Deguise *et al*. 2017). More recently, dysregulation of innate immunity was reported in pediatric SMA patients, as they exhibit treatment responsive changes in inflammatory cytokine profiles (Bonanno *et al*. 2022; Nuzzo *et al*. 2023). Accumulating evidence suggests that SMN loss disrupts the immune system, contributing to excessive neuroinflammation and neurodegeneration.

Here, we show that the transcriptomes and proteomes of SMA model flies similarly display evidence of dysregulated innate immunity. Specifically, these SMA models exhibited an increase in transcripts and proteins involved in the *Drosophila* Immune Deficiency (IMD) and Toll signaling pathways. Concordantly, these animals also frequently displayed pigmented nodules (a.k.a. melanotic masses) that correlated with the molecular signatures of activated immune signaling. Knockdown of specific downstream targets of these signaling pathways ameliorated formation of melanotic masses caused by *Smn* mutation or depletion. Overall, findings here suggest that SMN protein loss induces a hyperactivation of innate immune signaling and a melanization defense response that correlates with the phenotypic severity of SMA-causing missense alleles.

## METHODS

### *Drosophila* Strains and Husbandry

Fly stocks were maintained on molasses and agar at room temperature (25°C) in vials or half-pint bottles. As previously described, FLAG-*Smn^Tg^* transgenes were site-specifically integrated into a PhiC31 landing site (86Fb) that had been recombined into the *Smn^X7^* null background (Bischof *et al*. 2007; Praveen *et al*. 2012, 2014; Spring *et al*. 2019). The *Smn^X7^* null line was a gift of S. Artavanis-Tsakonis (Harvard University, Cambridge, USA). C15-GAL4 (Brusich *et al*. 2015) was a gift of A. Frank, University of Iowa (Iowa City, USA). All other GAL4/*UAS-RNAi* stocks were obtained from the Bloomington Drosophila Stock Center (BDSC), see Table S14 for details.

To generate larvae expressing a single *Smn* missense mutant allele, *Smn^X7^*/TM6B-GFP virgin females were crossed to *Smn^X7^*, *Smn^Tg^*/TM6B-GFP males at 25°C. To reduce stress from overpopulation and/or competition from heterozygous siblings, crosses were performed on molasses plates with yeast paste, and GFP negative (*Smn^X7^*, *Smn^Tg^*/*Smn^X7^*) larvae were sorted into vials containing molasses fly food during the second instar larval stage. Sorted larvae were raised at 25°C until the desired developmental stage was reached.

Experiments involving *UAS-Smn-RNAi* expression were carried out at 29°C to maximize expression from the GAL4/*UAS* system and, therefore, the degree of *Smn* knockdown. To maintain consistency across experiments, we used molasses plates with yeast paste and subsequent sorting for all *Smn-RNAi* experiments.

### Tandem Mass Tag (TMT) Sample Preparation

Cell lysates (100 µg; n=3) were lysed in 8M urea, 75 mM NaCl, 50 mM Tris, pH 8.5, reduced with 5mM DTT for 45 min at 37°C and alkylated with 15mM iodoacetamide for 30 min in the dark at room temperature. Samples were digested with LysC (Wako, 1:50 w/w) for 2 hr at 37°C, then diluted to 1M urea and digested with trypsin (Promega, 1:50 w/w) overnight at 37°C. The resulting peptide samples were acidified to 0.5% trifluoracetic acid, desalted using desalting spin columns (Thermo), and the eluates were dried via vacuum centrifugation. Peptide concentration was determined using Quantitative Colorimetric Peptide Assay (Pierce).

Samples were labeled with TMT10plex (Thermo Fisher). 40 µg of each sample was reconstituted with 50 mM HEPES pH 8.5, then individually labeled with 100 µg of TMT reagent for 1 hr at room temperature. Prior to quenching, the labeling efficiency was evaluated by LC-MS/MS (Liquid Chromatography and Tandem Mass Spectrometry) analysis of a pooled sample consisting of 1ul of each sample. After confirming >98% efficiency, samples were quenched with 50% hydroxylamine to a final concentration of 0.4%. Labeled peptide samples were combined 1:1, desalted using Thermo desalting spin column, and dried via vacuum centrifugation. The dried TMT-labeled sample was fractionated using high pH reversed phase HPLC (Mertins *et al*. 2018). Briefly, the samples were offline fractionated over a 90 min run, into 96 fractions by high pH reverse-phase HPLC (Agilent 1260) using an Agilent Zorbax 300 Extend-C18 column (3.5-µm, 4.6 × 250 mm) with mobile phase A containing 4.5 mM ammonium formate (pH 10) in 2% (vol/vol) LC-MS grade acetonitrile, and mobile phase B containing 4.5 mM ammonium formate (pH 10) in 90% (vol/vol) LC-MS grade acetonitrile. The ninety-six resulting fractions were then concatenated in a non-continuous manner into twenty-four fractions and dried down via vacuum centrifugation and stored at -80°C until further analysis.

### Liquid Chromatography-Tandem Mass Spectrometry (LC-MS/MS)

Twenty-four proteome fractions were analyzed by LC-MS/MS using an Easy nLC 1200 coupled to an Orbitrap Fusion Lumos Tribrid mass spectrometer (Thermo Scientific). Samples were injected onto an Easy Spray PepMap C18 column (75 μm id × 25 cm, 2 μm particle size) (Thermo Scientific) and separated over a 120 min method. The gradient for separation consisted of 5–42% mobile phase B at a 250 nl/min flow rate, where mobile phase A was 0.1% formic acid in water and mobile phase B consisted of 0.1% formic acid in 80% ACN.

For the proteome fractions, the Lumos was operated in SPS-MS3 mode (McAlister *et al*. 2014), with a 3s cycle time. Resolution for the precursor scan (m/z 350–2000) was set to 120,000 with a AGC target set to standard and a maximum injection time of 50 ms. MS2 scans consisted of CID normalized collision energy (NCE) 30; AGC target set to standard; maximum injection time of 50 ms; isolation window of 0.7 Da. Following MS2 acquisition, MS3 spectra were collected in SPS mode (10 scans per outcome); HCD set to 65; resolution set to 50,000; scan range set to 100-500; AGC target set to 200% with a 150 ms maximum inject time.

### TMT Data Analysis

TMT proteome RAW files were processed using Proteome Discoverer version 2.5. ‘TMT10’ was used as the quantitation method. Peak lists were searched against a reviewed Uniprot drosophila database (downloaded Feb 2020 containing 21,973 sequences), appended with a common contaminants database, using Sequest HT within Proteome Discoverer. Data were searched with up to two missed trypsin cleavage sites and fixed modifications were set to TMT peptide N-terminus and Lys and carbamidomethyl Cys. Dynamic modifications were set to N-terminal protein acetyl and oxidation Met. Quantitation was set to MS3, precursor mass tolerance was set to 10 ppm and fragment mass tolerance was set to 0.5 Da. Peptide false discovery rate was set to 1%. Reporter abundance based on intensity, SPS mass matches threshold set to 50, and razor and unique peptides were used for quantitation.

Statistical analysis was performed within Proteome Discoverer (version 2.4). Benjamini Hochberg corrected p-values (q-values) were calculated for each pairwise comparison, and statistical significance is defined as q-value<0.05. Log2 fold change (FC) ratios were calculated using the averaged normalized TMT intensities.

For Gene Ontology (GO) analysis, Uniprot protein IDs were converted to Flybase Gene IDs and gene symbols. GO enrichment was performed with FlyEnrichr, using the GO Biological Process (BP) category from AutoRIF (Chen *et al*. 2013; Kuleshov *et al*. 2016).

### RNA-seq Analysis

RNA-seq analysis was performed on fastq files retrieved from the NCBI Gene Expression Omnibus (GEO). GEO accesion numbers used here were: GSE49587, GSE81121, and GSE138183. Alignments of paired end reads were performed with HISAT2 and Ensemble release 109 of the *Drosophila melanogaster* genome (BDGP6.32) (Adams *et al*. 2000; Kim *et al*. 2019). Differential expression of transcripts was performed with kallisto and sleuth (Bray *et al*. 2016; Pimentel *et al*. 2017). For the determination of transcript abundance, the number of bootstrap samples was set at 100. StringTie and DESeq2 were used to determine differential gene expression (Love *et al*. 2014; Pertea *et al*. 2015, 2016).

### Scoring Melanotic Masses

Wandering third instar larvae were removed form vials, washed briefly in a room temperature water bath, dried, and placed on an agar plate under white light and 2X magnification. When melanotic masses were identified in a larva, both the size of the largest mass (size score) and the total number of masses (mass score) were qualitatively determined. Size scoring used the following criteria: small masses range in size from barely visible specks to smooth round dots with a diameter no more than 1/10th the width of the larva; medium masses range from anything larger than a small mass to those with a diameter up to 1/3 the larval width; large masses had a diameter greater than or equal to 1/3 the larval width. Larvae were manipulated to allow for observation of all sides/regions; observation was performed for at least 20 seconds in all cases.

### Statistical Analysis

GraphPad Prism version 7 was used to calculate *p*-values for comparison of melanotic masses, using a one-way ANOVA with a Dunnet correction for multiple comparisons.

### Data Availability

All *Drosophila* stocks are available upon request. The authors affirm that all data necessary for confirming the conclusions of the article are present within the article, figures, and tables. The tandem mass spectrometry labeling data have been deposited to the ProteomeXchange Consortium via the PRIDE partner repository (Perez-Riverol *et al*. 2022) using the dataset identifier PXD046801.

## RESULTS

### Quantitative proteomic analysis of *Smn* missense mutants identifies immune-induced peptides

Previously, we uncovered an increase in expression of genes associated with innate immunity in the transcriptomes of *Smn* null and missense mutant fly lines (Garcia *et al*. 2013, 2016); therefore, we sought to determine if the gene expression changes, identified by RNA-seq, are also reflected in the proteomes of hypomorphic *Smn* mutants. We therefore carried out proteomic analyses using tandem mass tag labeling and mass spectrometry (TMT-MS) on protein lysates from whole wandering third instar larvae. Animals expressing either Flag-*Smn* wild-type (WT) or SMA-causing missense mutant transgenes as their sole source of SMN protein were used. The transgenes were each inserted at the same ectopic locus and driven by the native *Smn* promoter in an otherwise *Smn^X7/X7^* null background (Praveen *et al*. 2012, 2014). We employed two different SMA patient-derived mutations located in distinct subdomains of the SMN protein, the Tudor domain (*Smn^Tg:V72G^*) and the tyrosine- and glycine-rich YG Box (*Smn^Tg:T205I^*), see Fig. 1A. The SMN Tudor domain binds symmetric dimethylarginine residues present at the C-termini of Sm proteins (Brahms *et al*. 2000, 2001), and the YG Box functions in SMN self-oligomerization (Liu *et al*. 1997; Talbot *et al*. 1997; Lorson *et al*. 1998; Martin *et al*. 2012; Gray *et al*. 2018; Gupta *et al*. 2021). As previously described, T205I is a Class 3 (semi-lethal, ∼10% eclosion) mutation, whereas V72G is more severe, categorized as a Class 2 mutation, as these animals all die as early pupae (Spring *et al*. 2019; Raimer *et al*. 2020).

**Figure 1.**
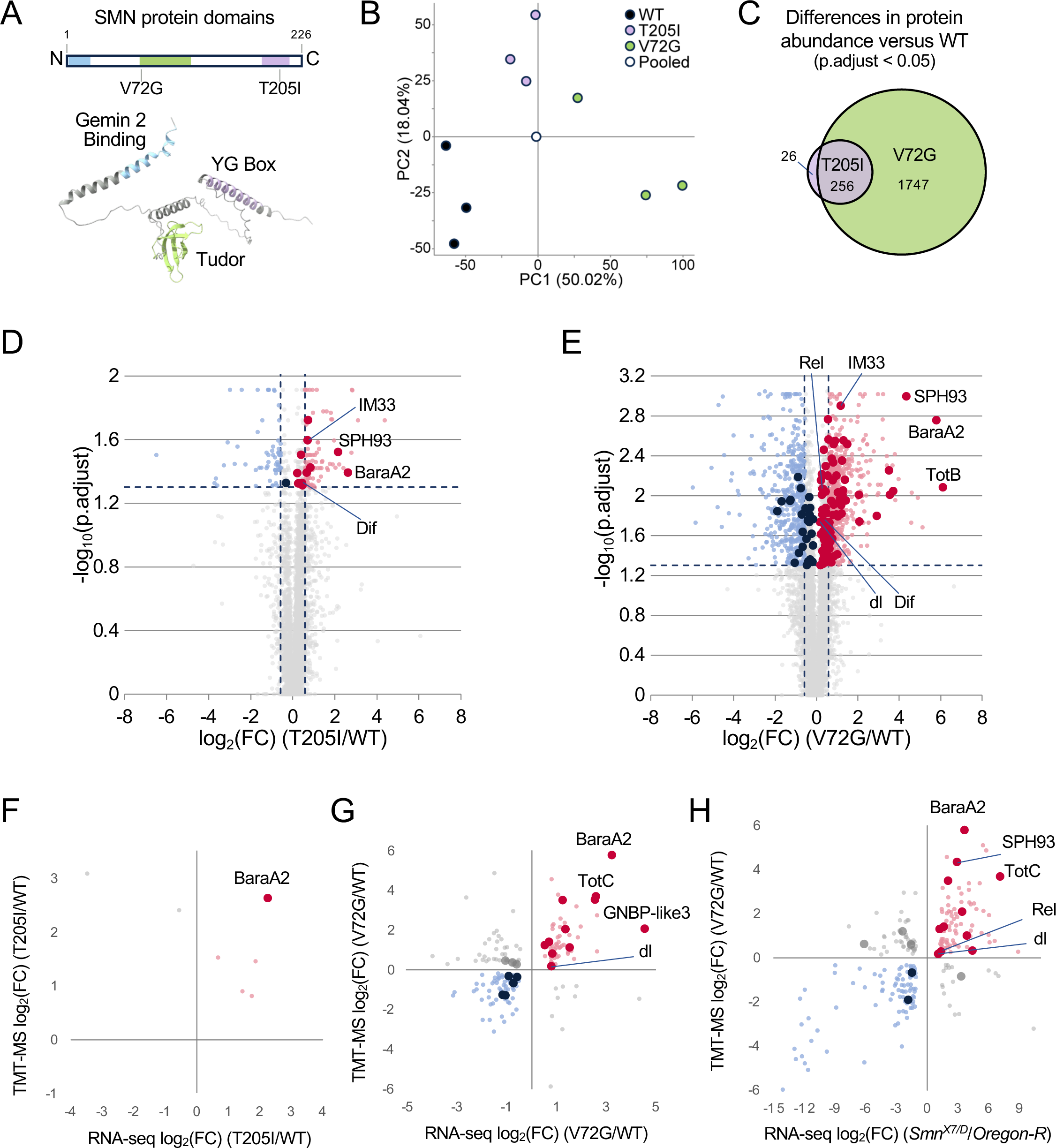
The proteomes and transcriptomes of *Drosophila Smn* hypomorphs display overlapping evidence for innate immune activation. **A)** A rectangular cartoon and an AlphaFold model of the relative positions of conserved domains of the *Drosophila* SMN protein and the location of the patient-derived missense mutations used here. **B)** Principal component analysis of total protein abundances in the *Smn* transgenic lines. *Smn* lines: WT (*Smn^Tg:WT;X7/X7^*); T205I (*Smn^Tg:T205I;X7/X7^*), Tyrosine (T) to Isoleucine (I); and V72G (*Smn^Tg:T205I;X7/X7^*), Valine (V) to Glycine (G). **C)** Venn diagram of overlapping protein differences in T205I and V72G relative to WT. **D)** Volcano plot of protein differences in the T205I line relative to WT. Proteins associated with innate immunity are indicated by larger dots. **E)** Volcano plot of protein differences in the V72G line relative to WT, and proteins associated with innate immunity are labeled as in (D). Dashed vertical bars in (D) and (E) indicate a Log2 FC ratio of +/- 0.58, and the horizontal dashed line corresponds to q-value = 0.05. **F)** Comparison of T205I proteome (y-axis) with T205I transcriptome (x-axis). The proteome and transcriptome are relative to the WT genotype. **G)** Comparison of V72G proteome (y-axis) with V72G transcriptome (x-axis). As in (F), the proteome and transcriptome are relative to WT. **H)** V72G proteome (y-axis) versus *Smn^X7/D^* null transcriptome (x-axis). The differential gene expression of the *Smn^X7/D^* transcriptome is relative to *Oregon-R*.

Overall, 5,857 *Drosophila* proteins were identified using TMT-MS (Tables S1-S3). Principal component analysis of TMT-MS quantified protein abundances showed good covariance levels (an average of ∼10% per sample) for the three different *Smn* transgenic lines we tested: *Smn^Tg:WT,X7/X7^* (WT), *Smn^Tg:V72G,X7/X7^* (V72G), and *Smn^Tg:T205I,X7/X7^* (T205I), see Fig. 1B. Among the proteins quantified, only 282 proteins were differentially expressed (p.adj <0.05, log_2_ fold change ±0.5) in the T205I mutant relative to WT control (Fig. 1C,D). Note that the control animals expressing the WT rescue transgene are known to be slightly hypomorphic to begin with (Praveen *et al*. 2014; Spring *et al*. 2019), so that may account for the small number of observed differences. In contrast, the V72G mutant exhibited 2,003 differentially expressed proteins relative to WT control (Fig. 1C,E). Most of the protein abundance differences found in the T205I mutant (90%) were also seen in the V72G mutant (Fig. 1C). The V72G and T205I hypomorphs each display significant defects (in viability, locomotion, etc.) relative to the WT controls, but the phenotype of the V72G animals is more severe than that of T205I (Praveen *et al*. 2014; Spring *et al*. 2019). Thus, the observed changes in protein abundance correlate with overall phenotypic severity (Fig. 1D-E).

### Immune dysregulation lies at the intersection of SMA model proteomes and transcriptomes

We took advantage of an early pupal RNA-seq dataset we had previously generated for *Smn* WT, T205I and V72G animals (Garcia et al. 2016) (Tables S4-S5) to carry out a multi-omic analysis of transcriptomes and proteomes. Although the TMT-MS experiment detected only a subset of the genes that can be analyzed by RNA-seq (e.g. 6,000 vs. 13,000), proteins that were significantly altered in the mutants also tended to display a similar trend on the transcriptome level. To this end, correlation plots of the log2 fold change ratios of the TMT-MS vs. total RNA-seq datasets showed good overall agreement between differences in RNA and protein abundance relative to the WT control (Fig. 1F,G).

Even though the milder T205I (Class 3) mutant had only ∼300 detectable changes at the protein level, and only seven overlapping RNA and protein changes (Fig. 1F), most of these (five out of seven) were increased in the T205I compared to WT). Notably, this includes the *Baramicin* locus (containing two identical genes, *BaraA1* and *BaraA2*) that encode an immune-induced antifungal peptide (Hanson *et al*. 2021; Hanson and Lemaitre 2022). For simplicity, we refer to all transcripts and proteins that mapped to this locus as *BaraA2 or* BaraA2, respectively (Fig. 1F). By comparison, the overlapping differences between the transcriptome and proteome of the more severe V72G (Class 2) mutant include increases in numerous immune-induced and stress responsive gene products (Fig. 1G). We note that analysis of the T205I transcriptome identified increases in many of these same immune-induced molecules that were not captured by TMT-MS (Table S4). Strikingly, we observed small but significant increases in core upstream signaling factors like the NF-kB ortholog dorsal (dl) and larger increases in defense-responsive and downstream stress-responsive targets like BaraA2, Turandot C (TotC), and Gram-negative bacteria binding-like protein 3 (GNBP-like3). Hence, our multi-omic approach further highlights the hyperactivation of innate-immune signaling that accompanies partial SMN loss-of-function.

For additional comparisons to the *Smn* missense mutant proteomes, we used polyA+- RNA-seq datasets from two different *Smn* null mutant lines (Li 2020; Garcia 2013) (Tables S6- S11). The *Smn^X7/D^*null mutant transcriptome identified an increase in *BaraA2* and *SPH93* (*Serine protease homolog 93*) transcripts in both T205I and V72G proteomes (Fig. 1H and Tables S1-S3). The overlap between the *Smn^X7/D^* transcriptome and the V72G proteome was even more remarkable and included the core NF-kB-like factor, Rel (Fig. 1H and Table S11). Thus, the overlapping differences between the *Smn* null and missense mutants suggest that the observed hyperactivation of immune signaling is a common feature of SMN loss.

A key strength of this multi-omic approach is the ability to detect mRNA and protein isoform-specific differences. For this analysis we employed an additional, probabilistic RNA-seq pipeline to quantify discrete mRNA isoforms and maintain pseudoalignment information from different splice junctions, but with a focus on differential expression of transcripts (Bray *et al*. 2016; Pimentel *et al*. 2017). Quantification of discernable transcript differences between *Smn* null and control animals revealed an increase in numerous transcripts associated with innate immunity in the mutants (Fig. S1A-B). Differences included changes in transcripts and proteins involved in innate immunity, such as the NF-kB orthologs dorsal (dl), Dorsal-related immunity factor (Dif), and Relish (Rel), (Fig. S1A-B).

Most striking, a comparison of the V72G proteome with the *Smn* null transcriptome revealed parallel isoform-specific changes for numerous transcripts and proteins (Fig. S1C). The congruous changes in RNA and protein isoforms included changes in molecules involved in innate immunity, including SPH93-RA/PA, TotC-RA/PA, GNBP-like3-RA/PA and Dif-RC/PC (Fig. S1C). In summary, the identification of overlapping changes in specific transcripts and protein isoforms further supports a uniquely well-coordinated activation of immune signaling in fly models of SMA.

### Partial loss of SMN function causes hyper-activation of innate immunity

SMA is a hypomorphic condition; total loss of function causes early developmental arrest and lethality (Schrank et al 1997; reviewed in O’Hern 2017). As detailed widely in the literature, *Smn* null mutants are thus poor disease models. Hence, we focused our efforts to identify drivers of the observed innate immune dysfunction on the *Smn* hypomorphs. Gene ontology (GO) analysis of protein abundance differences in the V72G dataset revealed a broad dysregulation of factors involved in pathogen defense response and innate immune signaling pathways (Fig. 2A and Tables S12-S13). These include proteins involved in melanization and humoral defense responses to bacterial, fungal, and viral pathogens (Figs. 2A-B). Although the V72G mutant exhibited numerous increases in proteins involved in defense response pathways, a few of these proteins were also significantly upregulated in the less severe T205I animals (Fig. 2B). Importantly, both mutants displayed small but significant increases in NF-κB transcription factors (Fig. 2B).

**Figure 2.**
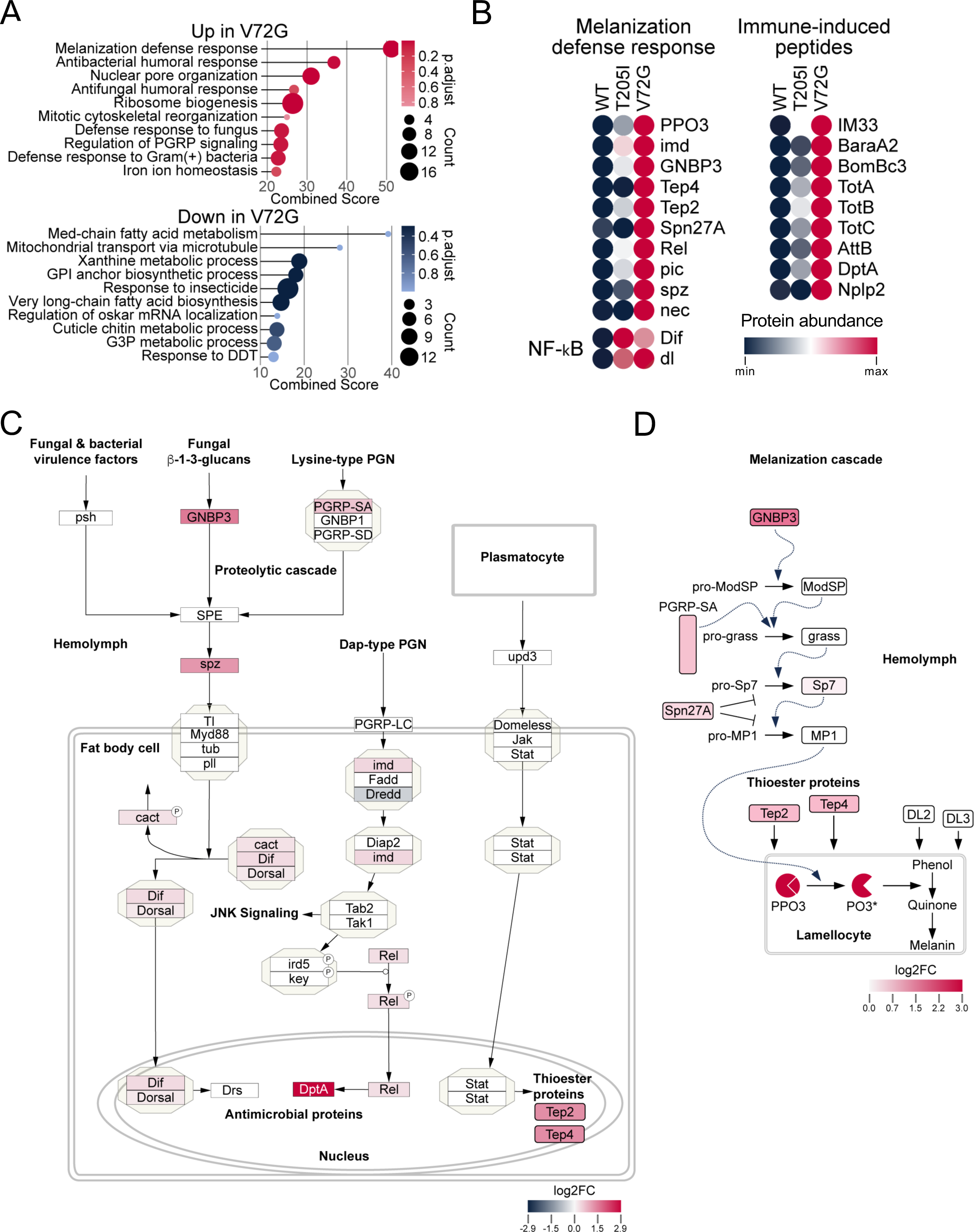
Proteins involved in *Drosophila* humoral and melanization defense responses are elevated in *Smn* mutant proteomes. **A)** Gene Ontology (GO) term analysis of protein differences in V72G. Adjusted p-values (p.adjust) and number of genes per GO term (Count) are shown at right, which is used to compute a combined score. **B)** Heat maps of select protein abundance differences from genes within the melanization defense response GO category, known immune-induced peptides, as well as for the NF-kB transcription factors Dorsal-related immunity factor (Dif) and dorsal (dl). **C-D**) Heatmap illustrations of TMT-MS data from V72G mutants. Log_2_-fold change (log2FC) values (mutant/control) for differentially expressed proteins are illustrated within the context of the Humoral Immune Response (Wikipathway WP3660, panel C) or the Melanization Defense Response (panel D) pathway and shaded according to the keys below each pathway.

Upregulation of defense response proteins occurs in the absence of an external immune challenge, supporting the notion that partial loss of SMN function causes hyper-activation of innate immune signaling. Consistent with this hypothesis, we frequently observed black, melanotic spots or granules in third instar *Smn* missense mutant larvae. Such granules are commonly referred to as pseudotumors, melanotic tumors, or melanotic masses (Minakhina and Steward 2006; Boulet *et al*. 2018). These structures typically form in response to pathogens, tissue damage, and necrosis, but this defense response can also be triggered by different genetic perturbations (Minakhina and Steward 2006; Williams 2007; Gold and Brückner 2015; Banerjee *et al*. 2019).

Irrespective of the trigger, melanotic masses often form in the larval hemolymph, and can be readily observed through the transparent body wall (Minakhina and Steward 2006). We therefore carried out a systematic analysis of larval melanization (Fig. 3) in a battery of ten hypomorphic, SMA-causing *Smn* missense alleles developed in our laboratory (Praveen et al. 2014; Spring et al. 2019). To quantify this phenotype, we scored both the size and number of melanotic masses in 50 wandering third instar larvae for each genotype. All lines examined displayed a statistically significant and robust increase in the presence of melanotic masses relative to the Oregon-R (OreR) controls (Fig. 3A). Larvae with a WT *Smn* transgene exhibited significantly fewer melanotic masses than *Smn* missense mutant lines but more than OreR (Fig. 3A), consistent with our previous observations that the Flag-*Smn* WT transgenic line is mildly hypomorphic (Praveen *et al*. 2014; Spring *et al*. 2019). Size scoring (Fig. 3B) and counts of the total number of melanotic masses per animal (Fig. 3C) show similar trends to the overall incidence of masses. The number of melanotic masses correlated with the previously characterized phenotypic severity of the different *Smn* missense mutations (Fig. 3A-D) (Spring *et al*. 2019). These observations suggest that the function of SMN in immune tissues is conserved from flies to mammals and that *Smn* mutations in the fly can be used to model peripheral defects of SMA in addition to the canonical SMA-related neuromuscular phenotypes.

**Figure 3.**
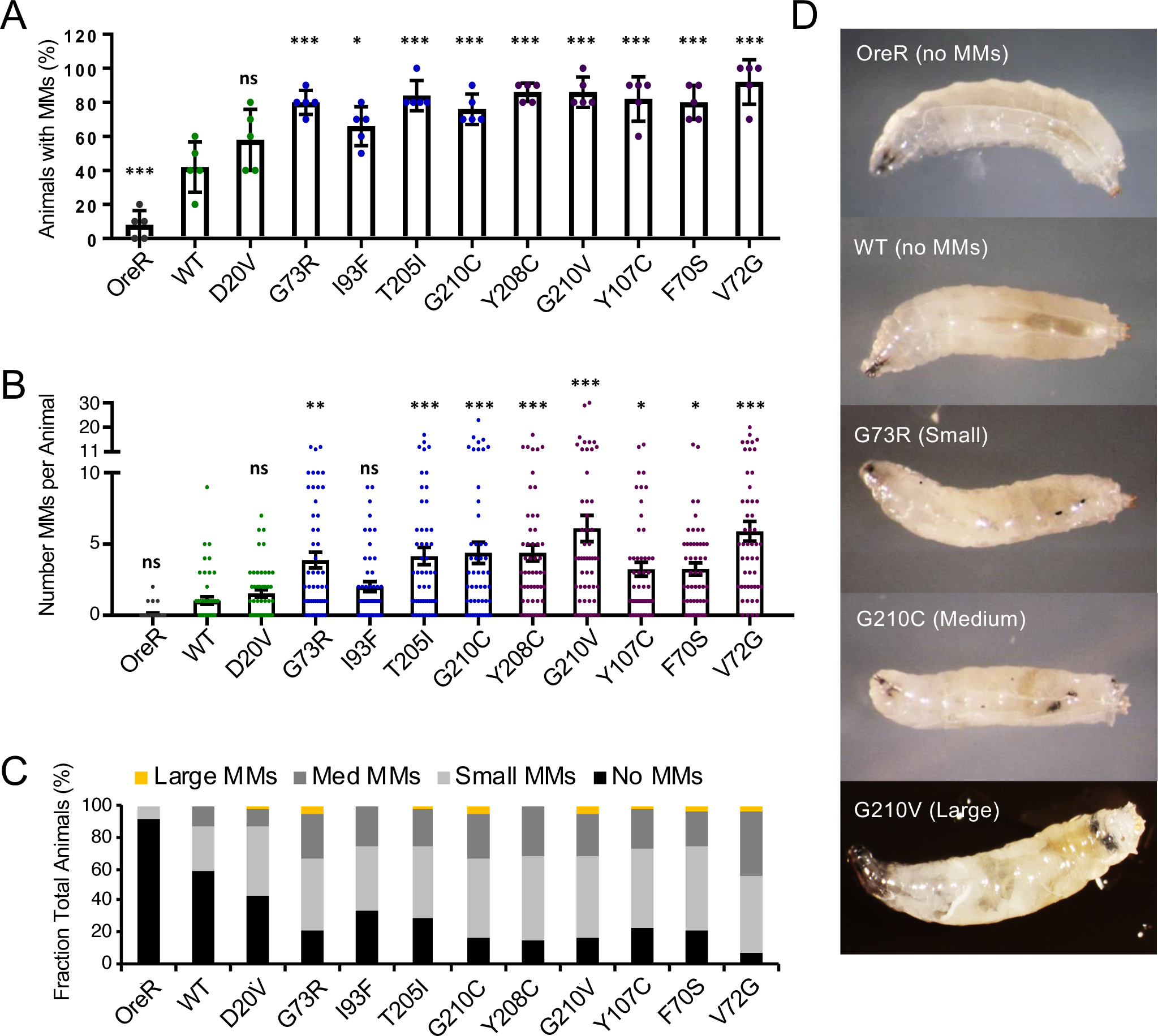
*Smn* missense mutants exhibit elevated melanotic masses. A-C) Melanotic mass (MM) data for wandering third instar larvae expressing *Smn* missense mutations. The data in each panel are a different measure of the melanotic mass phenotypes of the same set of larvae. **A)** Percent of larvae with one or more melanotic mass. Individual data points are the percent of larvae with MMs, 10 larvae per data point. **B)** The average number of melanotic masses per animal. Data points show the number of MMs in each animal. Number (N) = 50 larvae for each genotype. **C)** Qualitative size scoring of the largest melanotic mass in each larva. **D)** Representative images of melanotic masses in animals expressing *Smn* missense mutations. Bars show the mean, and error bars show standard error of the mean. Asterisks indicate *p*-values relative to WT: * < 0.05; ** < 0.01; and *** < 0.001.

### The SMN-dependent hyper-activation of melanization is tissue-specific

To determine if the melanotic masses in fly models of SMA are downstream effects of tissue specific SMN loss, we used the *Drosophila* GAL4/UAS system and RNA interference (RNAi) to deplete SMN in specific tissues (Perkins *et al*. 2015). We and others have previously employed this system to create partial SMN loss-of-function models that typically cause pupal lethality, although weakly viable adults can be obtained if the RNAi is performed at lower temperature, e.g. 25°C, see (Dimitriadi *et al*. 2010; Spring *et al*. 2019). Here, we employed two different *UAS:Smn* short hairpin (sh)RNA lines, P{TRiP.JF02057}attP2 (*Smn^JF^*-RNAi) and P{TRiP.HMC03832}attP40 (*Smn^HM^*-RNAi), at 29°C (Spring *et al*. 2019). Using a *daughterless* GAL4 driver (*da-Gal4*), we found that systemic SMN knockdown recapitulated the effects of the *Smn* missense mutations described above (Fig. 4A). Melanotic mass formation was dependent upon shRNA expression, as negative control lines (Gal4 driver-only, UAS:responder-only or OreR) showed no significant effects (Fig. 4A).

**Figure 4.**
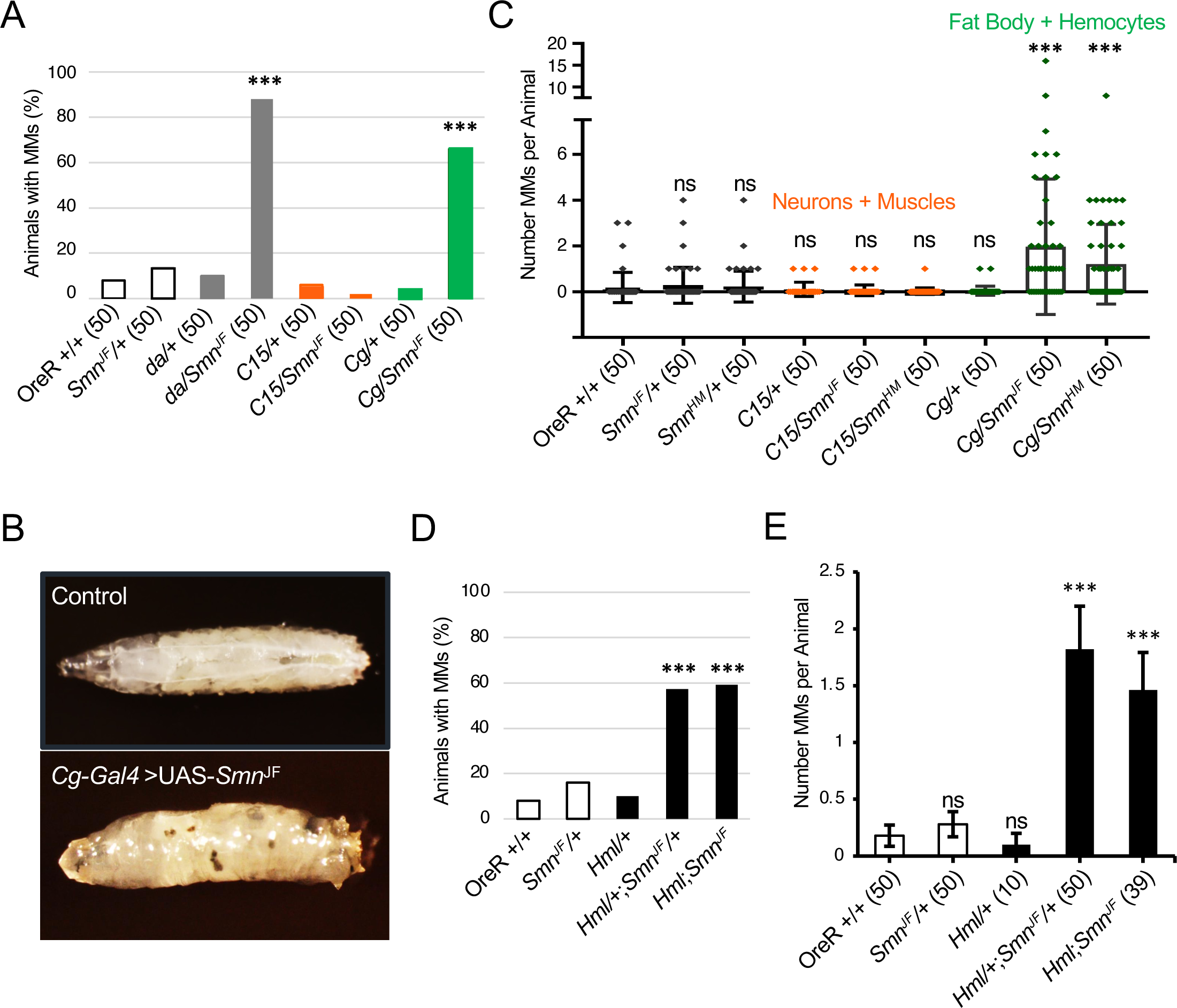
Targeted RNAi depletion of *Smn* in *Drosophila* immune cells yields melanotic masses and reduced viability. **A)** Fraction of larvae that display MMs. RNAi mediated knockdown of SMN was carried out using the *Drosophila* GAL4/UAS system to drive expression using UAS-*Smn^JF^* (P{TRiP.JF02057}attP2) together with the following GAL4 drivers: *da*, *daughterless* (*da*) for ubiquitous knockdown; *C15* (a composite driver that includes *elav-* (*embryonic lethal abnormal vision*), *sca- (scabrous*) and BG57-GAL4 see (Budnik *et al*. 1996; Brusich *et al*. 2015) for knockdown in both neurons and muscles; and *Cg (Collagen 4a1 gap)*, for knockdown in the fat body, hemocytes, and the larval lymph gland. OreR is the control strain. A plus sign (+) indicates a wild-type chromosome. **B)** Representative image of a wild-type control and MMs in a larva with SMN depleted in the fat body, hemocytes, and lymph gland (*Cg*- *GAL4*>*UAS*-*Smn^JF^*). **C)** Number of MMs per animal with and without SMN depletion, as in (A). *Smn^HM^* (P{TRiP.HMC03832}attP40) is an alternative UAS RNAi line that targets *Smn*. **D and E)** Fraction of larvae with MMs (D) and number of MMs per animal (E) using the hemocyte specific Gal4 driver *Hml (Hemolectin)* together with the *UAS-Smn^JF^* transgene. Control strains as per panel A.

In *Drosophila*, the immune response is coordinated by the fat body, an organ that is functionally analogous to the mammalian liver and adipose tissue (Hoffmann and Reichhart 2002; Ferrandon *et al*. 2007). The fat body signals to a group of macrophage-like cells, collectively called hemocytes (Banerjee *et al*. 2019). The molecular pathways and mechanisms that regulate hemocyte/macrophage development and activity are conserved from flies to humans (Williams 2007; Gold and Brückner 2015; Banerjee *et al*. 2019). When activated, hemocytes encapsulate invading particles and melanize them to sequester and kill pathogens (Banerjee *et al*. 2019). Depletion of SMN within the fat body and hemocytes (using *Cg-Gal4*) led to both a high frequency and number of melanotic masses per animal (Fig. 4A,B,C). In contrast, knockdown of SMN throughout the larval neuromusculature (using *C15-Gal4*) had no significant effect (Fig. 4A,C). Thus, the appearance of melanotic masses following depletion of SMN within immune cells rather than in neurons or muscles suggests that this phenotype is not a downstream consequence of neuromuscular dysfunction (Asha *et al*. 2003; Jenett *et al*. 2012; Hoffmann and Reichhart 2002; Ferrandon *et al*. 2007).

To ascertain whether melanotic mass formation was a consequence of SMN depletion within hemocytes, we carried out additional assays using the *Hemolectin-Gal4 (Hml-Gal4)* driver. As shown in Fig. 4D and 4E, knockdown of SMN specifically within hemocyte lineages also resulted in formation of larval melanotic masses. Therefore, we conclude that the observed melanization phenotype in response to SMN loss is derived (at least in part) from cell-autonomous defects in immune cells.

### Signaling pathways that regulate SMN-dependent melanization

To measure the relative contribution of various genes and pathways to the formation of melantoic masses induced by SMN knockdown, we next carried out a series of genetic modifier assays (Table S14). Given the results in Fig. 4, and the well-known function of the fat body in synthesizing and secreting antimicrobial peptides (AMPs) into the hemolymph (Hanson and Lemaitre 2020), we focused our screening efforts using the *Cg-Gal4* driver to reduce SMN levels by RNAi and then crossed in various mutations or secondary shRNA transgenes into this background.

The Toll, IMD and TNF (Tumor Necrosis Factor alpha, called Eiger in flies) signaling pathways use NF-kB transcription factors (dl, Dif and Rel) to turn on AMP genes (Hoffmann and Reichhart 2002; Lemaitre and Hoffmann 2007; Lindsay and Wasserman 2013; Hanson and Lemaitre 2020). Based on our multi-omic evidence (Figs. 1 and 2) showing overexpression of these NF-kB orthologs in our SMA models, we first ingressed heterozygous mutations for *dl* and *Rel* into the background of *Cg-Gal4/Smn^JF^*-RNAi flies to reduce dosage of these genes and then scored the resultant progeny for melanotic masses. As shown in Fig. 5A, mutants for *dl* and *Rel* suppressed the phenotype, reducing the average number of melanotic masses per larva. We also tested the dl/Dif regulatory factor, cactus. Contrary to our expectation, reduced dosage of cactus also reduced the number of melanotic masses. Mutations in *cactus* alone can cause melanotic masses (Minakhina and Steward 2006). However, since cactus levels are elevated in T205I and V72G animals (log2FC=0.26) and the well-documented autoregulatory feedback loop for this protein (Nicolas *et al*. 1998), the results of this modifier are inconclusive. Nevertheless, these data show that reducing gene dosage of downstream targets can suppress the melanization phenotype but throughput for this assay is quite low, often requiring generation of recombinants, and is limited by the genomic locations and availability of mutations of target genes.

**Figure 5.**
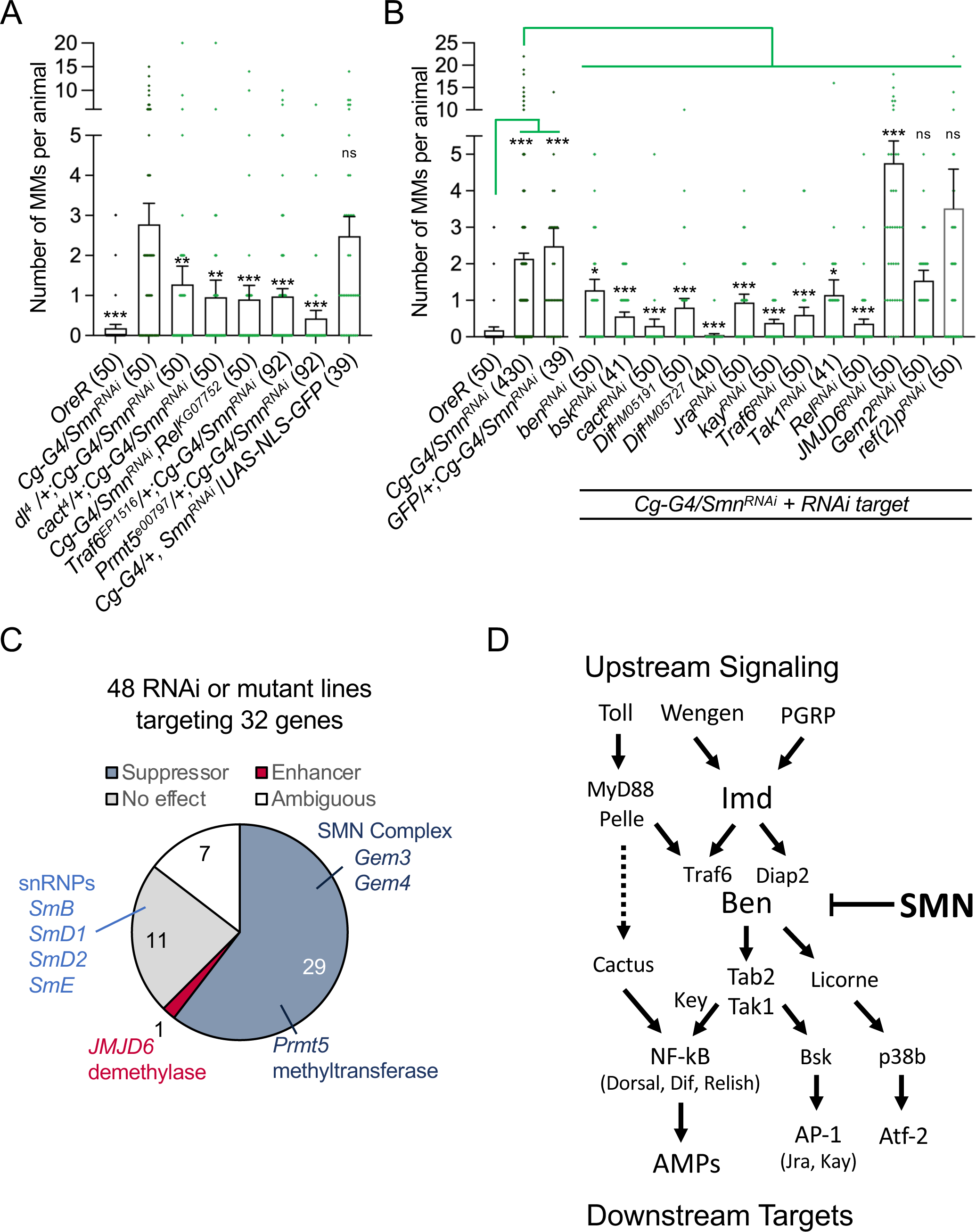
Innate immune signaling pathways contribute to MMs upon SMN depletion. **A)** Mutations in the IMD and Toll signaling pathways suppress the number of MMs per animal in *Smn* RNAi lines. Mutation of *Protein Arginine Methyltransferase 5* (*PRMT5*) also suppresses MMs upon depletion of SMN. **B)** Co-depletion of SMN and the indicated RNAi lines for members of the Toll and IMD pathways, *Jumonji domain containing 6* (*JMJD6*), *Gemin 2* (*Gem2*), and *refractory to sigma P* (*ref(2)P*). **C)** Pie chart of the identified enhancer and suppressors of MMs, resulting from *Smn* RNAi depletion in the fat body, hemocytes, and lymph gland. **D)** Model summarizing the role of SMN in the homeostatic regulation of the Toll, TNF and IMD signaling pathways in *Drosophila* larvae. Bendless (Ben) is an E2 ubiquitin conjugase that functions together with two different E3 ligases (Traf6 for Wengen/Toll and Diap2 for the PGRP pathway). The Immune Deficiency protein (Imd) serves not only as an upstream signaling factor, but also as a downstream target for K63-linked polyubiquitylation via Ben•Diap2. Ben thus sits at a node that connects several different signalling pathways and its activity is negatively regulated by SMN. Reduced levels of SMN thereby lead to hyperactivation of downstream targets.

To expand the scope of the investigation, we employed an RNAi-based candidate approach that couples *Cg-Gal4* mediated knockdown of *Smn* with the co-depletion of other factors. As a negative control for potential titration of GAL4 (which could reduce the efficacy of *Smn* knockdown) we co-expressed a UAS:NLS-GFP transgene. As shown in Fig. 5B, co-expression of a second UAS responder construct had no effect on the number of melanotic masses in the control larvae. In contrast, co-depletion of *Rel* gave similar results to those obtained with *Rel* mutants (compare Figs. 5A to 5B).

Next, we tested the effects of co-depleting SMN complex proteins and other known associated factors, see Table S14 for complete list. As shown, knockdown of snRNP components (SmB, SmD1, SmD2 and SmE) and one SMN complex member, Gemin2 (Gem2), had little effect on melanotic mass number (Figs. 5B,C). Co-depletion of two other SMN complex members, Gemin3 (Gem3; Shpargel et al. 2009), Gemin4/Gaulos (Gem4; Matera et al. 2019) and the arginine methyltransferase, Prmt5/dart5/capsuleen (Gonsalvez et al. 2006), suppressed the melanization phenotype (Fig. 5C). Interestingly, Gem3 and Gem4 were both previously shown to form complexes in S2 cells with the immune deficiency (imd) protein (Guruharsha *et al*. 2011), suggesting a potential role for Gemin subcomplexes in immune signaling. Prmt5 is a notable suppressor not only because knockdown of its corresponding arginine demethylase (JMJD6) enhanced the number of melanotic masses (Figs. 5B,C), but also because the Tudor domain of SMN is known to bind to dimethylated targets of Prmt5 (Friesen *et al*. 2001; Meister *et al*. 2001; Meister and Fischer 2002). We previously showed that complete loss of *Drosophila* Prmt5 function has little effect on snRNP assembly or organismal viability (Gonsalvez *et al*. 2008). Collectively, these data indicate that the presumptive SMN-interacting, innate immune signaling target of Prmt5 and JMJD6 is unlikely to be connected to SMN’s role in spliceosomal snRNP biogenesis. We therefore sought to test other candidate signaling factors that interact with SMN.

A common feature of the Toll (Toll), IMD (PGRP) and TNF/Eiger (Wengen) signaling pathways (Fig. 5D) is a protein complex that forms a platform for K63-linked ubiquitylation and recruitment of downstream factors like Tak1 (TGF-β activated kinase 1), Tab2 (TAK1-associated binding protein 2), and key (kenny, a.k.a. NEMO). Analogous complexes function within the mammalian TLR (Toll like receptor) and TNFa (Tumor necrosis factor alpha) signaling cascades (Shen *et al*. 2001; Cha *et al*. 2003; Ma *et al*. 2014; Ding *et al*. 2022). In mammals, TLR signaling involves the E3 ligase Traf6 (TNF Receptor Associated Factor 6), whereas TNFa signaling utilizes Traf2 (Igaki and Miura 2014; Ma *et al*. 2014; Sharma *et al*. 2021). In flies, a single protein, called Traf6/dTRAF2 can perform both functions (Shen *et al*. 2001; Kauppila *et al*. 2003). As in humans, fly Traf6 likely interacts with the E2 conjugating enzyme Ubc13/bendless (Ma *et al*. 2014). Ubc13/bendless and the Ubiquitin-conjugating enzyme variant 1A (Uev1A) activate Tak1, a downstream kinase in the IMD pathway, although Traf6 appears to be dispensible for this activation, at least in S2 cells (Zhou *et al*. 2005; Chen *et al*. 2017).

Intriguingly, human TRAF6 was shown to co-precipitate with SMN (Kim and Choi 2017). The authors hypothesized that SMN might serve as an negative regulator of NF-kB signaling, although the effect could be indirect (Kim and Choi 2017). We therefore tested this idea *in vitro* with purified components, and found that human GST-TRAF6 interacts directly with SMN•Gem2 heterodimer (Fig. S2A). Experiments aimed at determining if this biochemical interaction was conserved in the fly were inconclusively negative. Transgenic overexpression of Flag-tagged fruit fly Traf6 (tub-Gal4 > UAS:Flag-Traf6) failed to co-immunoprecipitate endogenous SMN (Fig. S2B). As measured by western blotting, we also failed to detect *Drosophila* Traf6 in co-precipitates from embryonic lysates expressing Flag-SMN as the sole source of SMN protein. However, our previous AP-MS (affinity purification followed by mass spectrometry) analysis of Flag-SMN pulldowns identified the E2 conjugase and Traf6 binding partner, Ubc13/ben (Gray *et al*. 2018).

Given that biologically important interactions are not necessarily biochemically stable enough to withstand a pulldown assay, we decided to test ben and Traf6 by genetic interaction in the larval melanization assay. As shown in Figs. 5A and 5B, reduction in dosage of either *Traf6* or *ben* resulted in a significant decrease in the number of melanotic masses per animal, compared to that of the SMN RNAi-only control. In summary, these observations show that Toll, IMD, and TNF-Eiger signaling pathways are disrupted following loss of SMN expression within the immune system (fat body and hemocytes), leading to formation of melanotic masses in fly models of SMA.

## DISCUSSION

Our multi-omic investigation of fly models of SMA supports a role for dysregulated innate immunity in the peripheral pathophysiology associated with the disease in humans. The molecular signatures of an activated immune response were readily apparent in the whole-animal transcriptomes and proteomes of two hypomorphic *Smn* mutants. Moreover, we observed aberrant immune activation in all SMA models examined, including very mild models (Fig. 3) that do not display neuromuscular or viability defects in the larval stage (Spring *et al*. 2019). Furthermore, the degree of immune activation, as measured by larval melanotic mass formation (Fig. 3) correlated well with phenotypic class of the mutations (Spring *et al*. 2019).

That is, Class 2 SMA alleles had the most melanotic masses, Class 4 the fewest, and Class 3 had an intermediate number (Fig. 3B). These results are notably consistent with recent findings of immune dysregulation in mammalian models of SMA and in pediatric SMA patients (Zhang *et al*. 2013; Deguise and Kothary 2017; Khairallah *et al*. 2017; Thomson *et al*. 2017; Deguise *et al*. 2017; Vukojicic *et al*. 2019; Bonanno *et al*. 2022; Nuzzo *et al*. 2023). Furthermore, our work suggests that this conserved dysregulation of innate signaling is a primary effect of SMN loss in immune cells and tissues rather than a secondary consequence of SMN loss elsewhere. The extent to which the dysregulation of immune systems contributes to neuroinflammation and neuromuscular degeneration in SMA remains to be determined.

### Neurodegeneration and the sustained activation of innate immunity

Emerging evidence suggests that a hyperactivation of innate immunity is a common feature of neurodegeneration. Our finding that downstream targets of NF-kB proteins are upregulated in *Smn* hypomorphs is particularly revealing. Overexpression of NF-kB and other innate immune targets via the cGAS-STING (cyclic GMP-AMP Synthase-Stimulator of Interferon Response Genes) pathway contributes to disease progression in a mouse model of Alzheimer’s disease (Xie *et al*. 2023). NF-kB-related signaling has been implicated in the pathogenesis of ALS (Amyotrophic Lateral Sclerosis), also likely involving Toll-like receptors and the cGAS-STING pathway (Swarup *et al*. 2011; Egawa *et al*. 2012; Zhao *et al*. 2015; Yu *et al*. 2020; Lee and Woodruff 2021). In agreement with these findings, *Drosophila* models of Alzheimer’s disease, ALS, Ataxia-telangiectasia, and retinal degeneration further suggest that sustained activation of innate immunity contributes to neurodegeneration (Tan *et al*. 2008; Chinchore *et al*. 2012; Petersen and Wassarman 2012; Petersen *et al*. 2012; Han *et al*. 2020). During development, *Drosophila* orthologs of the cGAS-STING pathway function to condition the innate immune system, but the consequences of sustained activation of this pathway and the potential contribution to neurodegeneration remain to be determined (Cai *et al*. 2022; Wang *et al*. 2022).

In addition to large effects on innate immune signaling, our combinatorial, multi-omic approach provides insight into more subtle molecular consequences of SMN loss. For example, the proteome of the V72G mutant displayed altered protein levels for several SMA disease modifiers: CG17931/Serf, coronin (coro), and Zinc finger protein 1 (Zpr1), see Table S1 (Scharf *et al*. 1998; Ahmad *et al*. 2012; Hosseinibarkooie *et al*. 2016; Wirth *et al*. 2017; Zhuri *et al*. 2022). The reduction in levels of CG17931, an ortholog of Small EDRK-Rich Factor 1A (SERF1A/H4F5), are consistent with earlier findings in a different fly *Smn* mutant (Ghosh 2017). Among these three putative modifiers, Zpr1 is notable for its reported protein-protein interaction with SMN and snRNP import proteins (Gangwani *et al*. 2001; Narayanan *et al*. 2002). In addition, the latter two factors have been implicated in R-loop resolution and subsequent DNA-damage response (Zhao *et al*. 2016; Kannan *et al*. 2019; Cuartas and Gangwani 2022). The proteomes of both the mild T205I and severe V72G show evidence of a DNA-damage response (Table S1). In an unrelated study, cytoplasmic R-loop accumulation and DNA-damage response were recently linked to the activation of innate immunity via the Toll-like receptor and the cGAS-STING signaling pathways (Crossley *et al*. 2023). Future studies will be required to test the correlation, noted here, between the proteomic signatures of a DNA-damage response and the activation of innate immunity in these hypomorphic SMA models.

### SMN, K63-linked polyubiquitylation and immune signaling networks

In mammals and flies, the TLR/Toll and TNF/IMD signaling pathways function through analogous enzymatic cascades and complexes. Prominently featured in these pathways are receptor-proximal adaptor proteins (e.g. RIP1/Imd) that are activated by K63-linked ubiquitylation (K63-Ub) (Valanne *et al*. 2011; Lindsay and Wasserman 2013; Kietz and Meinander 2023). The protein complex that carries out these crucial post-translational modifications includes the E2 conjugating enzymes and cofactors Ubc13/bendless (Ben), Uev1a, and Ubc5/effete, along with two other RING-domain E3 ligases, Diap2 or Traf6 (see Fig. 5D). The presence of these K63-Ub oligomers triggers binding of Tab2 and key, leading to activation of the downstream kinase Tak1. Traf6 likely plays both enzymatic and structural roles in this process (Strickson *et al*. 2017).

Tak1 phosphorylation of I-kappaB kinase, mediated by binding Tab2 and key, leads to translocation of NF-kB transcription factors to the nucleus, and expression of antimicrobial peptide (AMP) genes (Fig. 5D). Traf6, Diap2 and Ben thus constitute an evolutionarily conserved node or nexus through which multiple intracellular signaling pathways are connected (Fig. 5D). The work here identifies SMN as a negative regulator of this complex, supported by both biochemical (Fig. S2A and (Kim and Choi 2017; Gray *et al*. 2018)) and genetic (Figs. 5A-B) interactions. In summary, we show that partial loss of SMN function (either by mutation or depletion) results in the sustained activation of innate immunity.

Our proteomic analyses of mild and intermediate fly models of SMA reveal clear signatures of an immune response in the absence of an external challenge. These include, but are not limited to, overexpression of AMPs (Figs. 1 and 2). Notably, Ganetzky and colleagues have shown that ectopic expression of individual AMP genes can bypass this immune signaling cascade and cause disease, as the neural overexpression of AMP transgenes is sufficient to cause neurodegeneration in the fly brain (Cao *et al*. 2013). Although the precise mechanisms remain unclear, neuroinflammatory responses like those identified here are likely to contribute to the pathophysiology of neurodegenerative diseases like Spinal Muscular Atrophy.

---

**Supplemental Figure S1.**
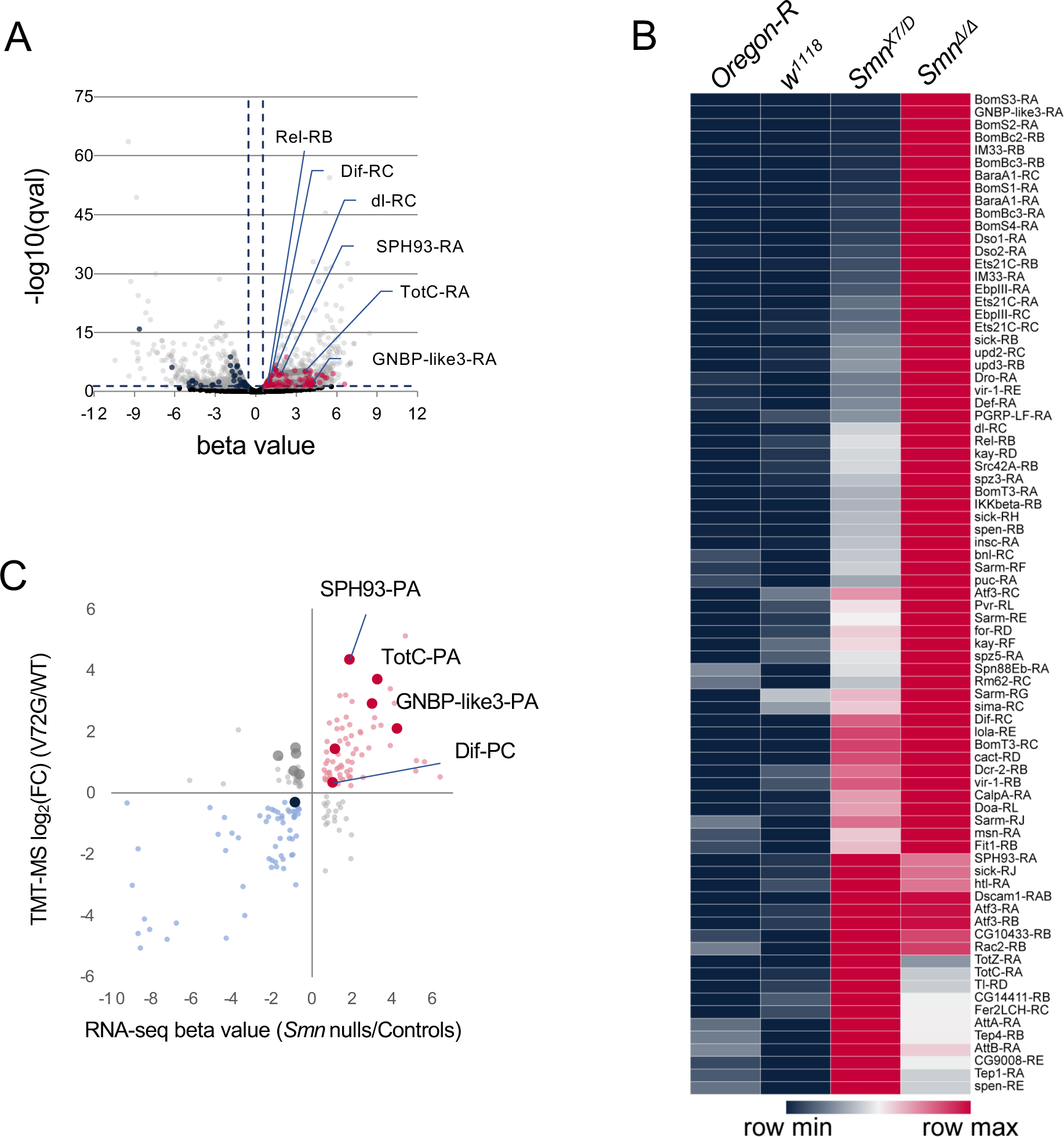
Isoform-specific differences in *Smn* mutants versus controls. **A)** Volcano plot of differentially expressed transcripts in *Smn* null animals. Transcripts associated with innate immunity are indicated with red circles and a subset of those are labeled with transcript symbols for the specific mRNA isoform difference. The axis shown: a Benjamini-Hochberg (False Discovery Rate (FDR) < 0.05) adjusted *p*-value (qval) and a Wald test-derived representation of a normalized fold change (beta factor). **B)** The heat map displays the respective mean transcripts per million reads for the different genotypes used in (A). The values are scaled and normalized per row (z-score). The heat map shows approximately half of the differentially expressed transcripts from (A). **C)** Scatter plot comparison of isoform-specific protein changes identified in the V72G proteome versus isoform-specific RNA changes found in the *Smn* null transcriptome. RNA and proteins associated with innate immunity are represented with bigger dots and labeled.

**Supplemental Figure S2.**
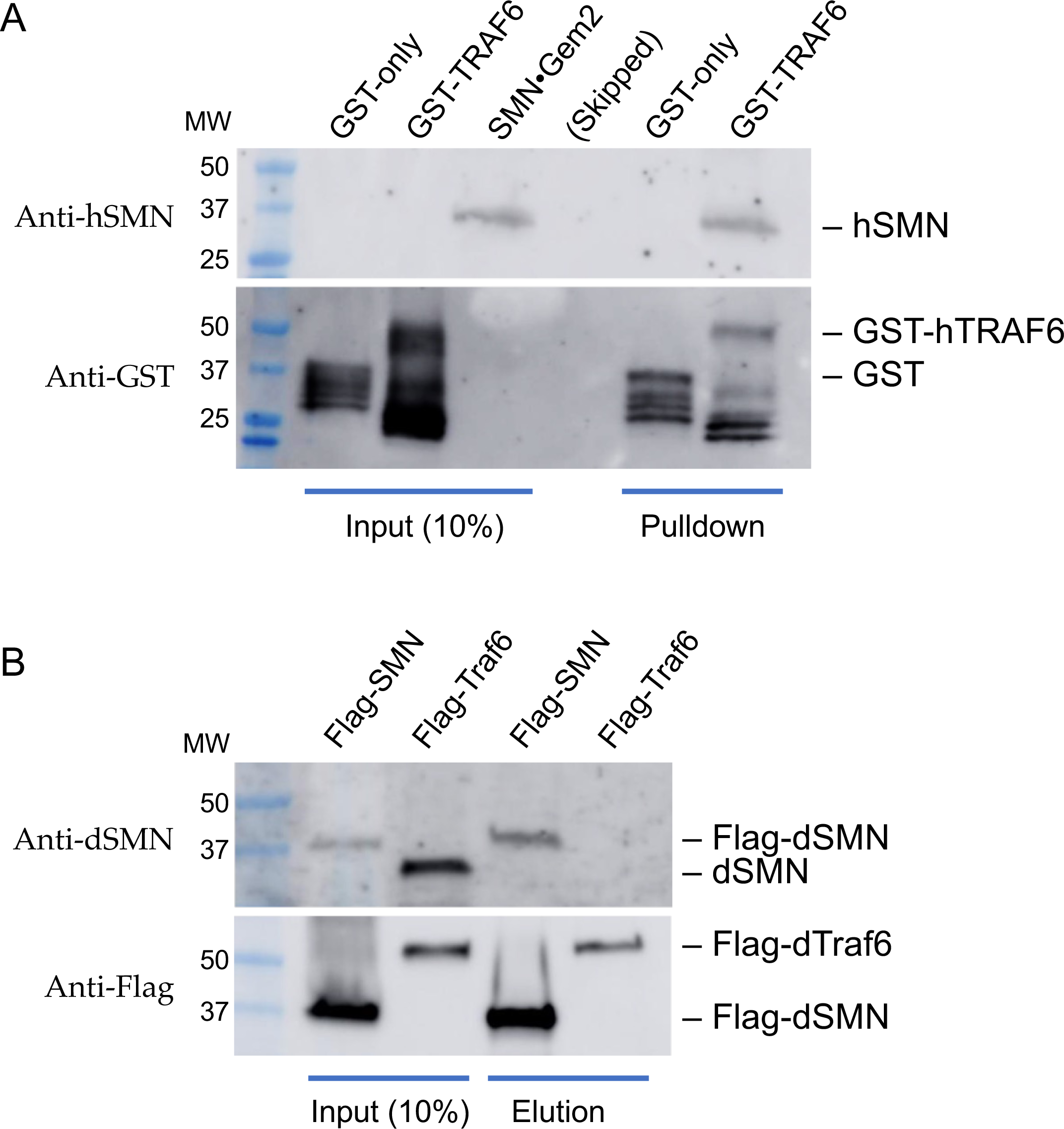
Evaluation of protein-protein interactions. **A)** GST-pulldown experiment using recombinant human GST-TRAF6 and SMN•Gem2. GST and GST-TRAF6 were expressed in E.coli and purified using anti-Glutathione beads. Pulldown assays were performed and analyzed by western blotting with either anti-hSMN (top) or anti-GST (bottom) antibodies. As shown, GST-hTRAF6 interacts directly with human SMN•Gem2. **B)** Flag-pulldown experiment using lysates from tub-Gal4 > UAS:Flag-dTraf6 animals (Flag-Traf6) or from control animals bearing a *Flag-Smn* transgene (Gray et al. 2018) as the only source of SMN protein (Flag-SMN). Inputs are on the left and proteins eluted from the Flag beads following pulldowns are on the right. As shown, Flag-SMN co-purifies with itself in the control lysates but Flag-Traf6 fails to pull down endogenous dSMN in the experimental cross.

